# EXoO-Tn: Tag-n-Map the Tn Antigen in the Human Proteome

**DOI:** 10.1101/840298

**Authors:** Weiming Yang, Minghui Ao, Angellina Song, Yuanwei Xu, Hui Zhang

**Affiliations:** Department of Pathology, Johns Hopkins University School of Medicine, Baltimore, Maryland, USA

**Keywords:** Tn antigen, site-specific, O-GalNAc, glycoproteomics, cancer

## Abstract

Tn antigen (Tn), a single N-acetylgalactosamine (GalNAc) monosaccharide attached to protein Ser/Thr residues, is found on most solid tumors yet rarely detected in adult tissues, featuring it one of the most distinctive signatures of cancers. Although it is prevalent in cancers, Tn-glycosylation sites are not entirely clear owing to the lack of suitable technology. Knowing the Tn-glycosylation sites will spur the development of new vaccines, diagnostics, and therapeutics of cancers. Here, we report a novel technology named EXoO-Tn for large-scale mapping of Tn-glycosylation sites. EXoO-Tn utilizes glycosyltransferase C1GalT1 and isotopically-labeled UDP-Gal(^13^C_6_) to tag and convert Tn to Gal(^13^C_6_)-Tn, which has a unique mass being distinguishable to other glycans. This exquisite Gal(^13^C_6_)-Tn structure is recognized by OpeRATOR that specifically cleaves N-termini of the Gal(^13^C_6_)-Tn-occupied Ser/Thr residues to yield site-containing glycopeptides. The use of EXoO-Tn mapped 947 Tn-glycosylation sites from 480 glycoproteins in Jurkat cells. Given the importance of Tn in diseases, EXoO-Tn is anticipated to have broad utility in clinical studies.

## Introduction

Over decades of biomedical investigations, it was found that one of the most distinctive features of cancers is the expression of Tn antigen (Tn), which is an N-acetylgalactosamine (GalNAc) attached to protein Ser/Thr residues via an O-linked glycosidic linkage ^1^. A variant of Tn is STn, which has an addition of sialic acid monosaccharide ^1^. Tn establishes its nature as a pan-carcinoma antigen by finding of its expression in 10-90% of the solid tumor including lung, prostate, breast, colon, pancreas, gastric, stomach, ovary, cervix, bladder ^1-3^. In sharp contrast, the expression of Tn in adult tissue is rare ^4^, making it an attractive target for anti-cancer applications. For instance, Slovin et al. report a Phase I clinical trial using a vaccine consisting of synthetic Tn on a carrier protein for prostate cancer ^5^. Studies explore the potential of Tn for early diagnostics ^6-8^ and prognostics of cancers ^9-11^. To treat cancers, Posey et al. report the development of engineered CAR-T cells that target Tn on mucin protein MUC1 (MUC1-Tn) for killing cancer cells ^12^. Also, a Phase I clinical trial using MUC1-Tn specific CAR-T cells started for treating patients with head and neck cancer ^13, 14^. Despite a noteworthy link between Tn and cancers, the underlying mechanism causing the expression of Tn in cancers is not entirely clear. It may involve glycosyltransferase C1GalT1 and its chaperone C1GalT1C1 also called Cosmc ^15^. Defective mutation in Cosmc is reported to affect the function of C1GalT1 for elongating Tn to normal O-glycan structures ^15, 16^. Furthermore, Tn is involved in IgA nephropathy (IgAN, also known as Berger’s disease) that is the most common glomerular disease in the world ^3, 17, 18^. A large percentage of patients with IgAN progress to kidney failure, also called end-stage renal disease (ESRD) ^3, 17^. The cause of IgAN may involve the expression of Tn and STn on hinge region of IgA1 ^3^.

Although Tn is structurally simple, identification of its glycosylation sites and the carrier proteins in the complex samples is highly challenging due to the lack of suitable technology. Limited information regarding Tn-glycosylation sites and carrier proteins hamper the understanding of the role of Tn in cancer biology and the development of new strategies targeting cancers. Current methods for mapping Tn-glycosylation sites include the use of VVA lectin or hydrazide chemistry for the enrichment of Tn-glycopeptides, followed by LC-MS/MS for site localization ^19, 20^. Jurkat T cells expressing Tn and STn, due to the mutation in Cosmc, are often used as a model system to evaluate the effectiveness of methods. Using VVA lectin chromatography and ETD-MS2, Steentoft et al. identify 68 O-glycoproteins in Jurkat cells ^19^. Zheng et al. use galactose oxidase to oxidize Tn followed by solid-phase capture using hydrazide chemistry and release of Tn-glycopeptides using methoxyamine ^20^. Subsequent analysis using HCD-MS2 identifies 96 O-glycoproteins in three experiments with 87 glycosylation sites being localized in the first experiment of Jurkat cells ^20^. We, however, anticipate that about a thousand Tn-glycosylation sites remain to be mapped in Jurkat cells because 1,295 O-linked glycosylation sites are mapped in CEM cells, a human T cell line, using a method named EXoO developed in previous study ^21^. It appears that the development of a technology capable of large-scale mapping of Tn-glycosylation sites would be a significant advance in technology and cancer biology.

Here, we introduce a new technology named EXoO-Tn that tags Tn and maps its glycosylation sites in a large-scale. EXoO-Tn utilizes two highly specific enzymes in a one-pot reaction for concurrent tagging of Tn and mapping of its glycosylation sites. The first enzyme is glycosyltransferase C1GalT1, which catalyzes UDP-Gal to add a galactose to Tn. When isotopically-labeled UDP-Gal(^13^C_6_) is used, Gal(^13^C_6_)-Tn is formed. The Gal(^13^C_6_)-Tn has a unique mass tag distinguishable to endogenous Gal-GalNAc and other glycans. The second enzyme is an endoprotease named OpeRATOR, which cleaves at N-termini of Ser/Thr residues occupied by the Gal(^13^C_6_)-Tn to release site-containing Gal(^13^C_6_)-Tn-glycopeptides with the glycosylation sites positioning at the N-termini of peptide sequences. The two enzymes are synergistically integrated with the use of solid-phase for optimal removal of contaminants and efficient isolation of site-containing Gal(^13^C_6_)-Tn-glycopeptides. A Proof-of-principle of EXoO-Tn was developed using a synthetic Tn-glycopeptide. The performance of EXoO-Tn was evaluated using Jurkat cells.

## Results

### Principle of EXoO-Tn

EXoO-Tn includes six steps (Fig. 1). (i) Digestion: proteins extracted from samples are digested to peptides. Amino groups on the side chain of Lys residues are modified using guanidination on C18 cartridge. (ii) Enrichment: Tn-glycopeptides are enriched using VVA lectin. (iii) Conjugation: the enriched glycopeptides are conjugated to aldehyde-functionalized solid-phase through amino groups at the peptide N-termini. (iv) Tn-engineering: Tn is catalyzed to Gal(^13^C_6_)-Tn using C1GalT1/C1GalT1C1 and UDP-Gal(^13^C_6_). C1GalT1/C1GalT1C1 is specific to modify Tn. The Gal(^13^C_6_)-Tn has a unique mass that is distinguishable to endogenous Gal-GalNAc and other glycans in the samples. (v) Release: site-containing Gal(^13^C_6_)-Tn-glycopeptides are specifically released from solid-phase using OpeRATOR enzyme, which cleaves N-termini of Gal(^13^C_6_)-Tn-occupied Ser/Thr residues. (vi) Analysis: the released glycopeptides are analyzed using LC-MS/MS and software tools.

**Figure 1.**
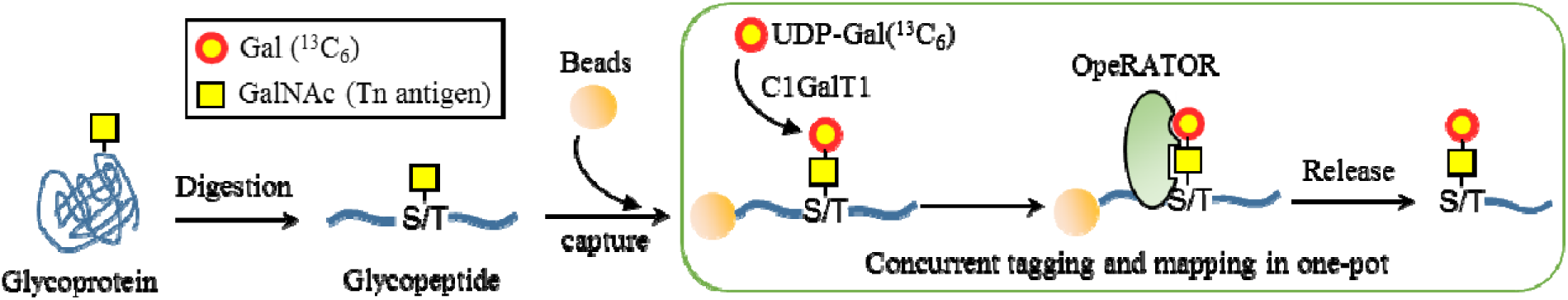
Strategy of EXoO-Tn for tagging of Tn and mapping its glycosylation site.

To show the feasibility of EXoO-Tn, a synthetic Tn-glycopeptide VPSTPPTPS(α-GalNAc)PSTPPTPSPSC-NH2 was used (Fig. 2A top left panel). The use of C1GalT1 and UDP-Gal converted Tn to Gal-Tn produced a charge +2 Gal-Tn-glycopeptide at 1149.54 *m/z* (Fig. 2A top middle panel), an increase of ∼162 Da corresponding to the mass of a galactose compared to its unmodified counterpart at 1068.51 *m/z* (Fig. 2A top left panel). The Gal-Tn-glycopeptide could be digested by OpeRATOR to yield site-containing glycopeptide S(Gal-Tn)PSTPPTPSPSC-NH2 at 761.34 *m/z* and peptide VPSTPPTP at 795.42 *m/z* (Fig. 2A bottom middle panel). To distinguish the newly engineered Gal-Tn from endogenous Gal-GalNAc and other glycans, the UDP-Gal was substituted by an isotopically-labeled UDP-Gal(^13^C_6_). The Gal(^13^C_6_) has all six carbon molecules in galactose labeled with carbon-13 featuring an increment mass of 6 Da. The use of C1GalT1 and UDP-Gal(^13^C_6_) successfully converted Tn to Gal(^13^C_6_)-Tn with a unique mass tag of 371 and yielded a charge +2 Gal(^13^C_6_)-Tn-glycopeptide at 1152.55 *m/z* (Fig. 2A top right panel), which had an increase of ∼6 Da compared to its charge +2 Gal-Tn counterpart at 1149.54 *m/z* (Fig. 2A top middle panel). The site-containing glycopeptide S(Gal(^13^C_6_)-Tn)PSTPPTPSPSC-NH2 and peptide VPSTPPTP at 764.35 and 795.42 *m/z*, respectively, was generated after OpeRATOR digestion (Fig. 2A bottom right panel). The Gal(^13^C_6_)-Tn-glycopeptide had an increase of ∼6 Da compared to its Gal-Tn or endogenous Gal-GalNAc counterpart at 761.34 *m/z* (Fig. 2A bottom middle panel). Next, the MS/MS spectra of site-containing Gal(^13^C_6_)-Tn-glycopeptides were analyzed using HCD-MS2 to identify spectral feature for improvement of confidence of identification. As an illustration, an MS/MS spectrum of site-containing Gal(^13^C_6_)-Tn-glycopeptide from analysis of Jurkat cells was shown (Fig. 2B). A diagnostic oxonium ion generated by HCD fragmentation was observed at 372 *m/z* for the Gal(^13^C_6_)-Tn (Fig. 2B). The presence of the diagnostic oxonium ion at 372 *m/z* was utilized in the data interpretation. The Gal(^13^C_6_)-Tn-glycosylation site was informed to be the Thr residue at the N-terminus of the identified peptide sequence (Fig. 2B). Other fragmentation ions in the MS/MS spectrum, including oxonium ions, peptide b- and y-ions, and peptide ion supported the identification of the glycopeptide (Fig. 2B). The analysis of glycopeptides demonstrated the key enzymatic steps in EXoO-Tn to distinguish Tn from Gal-GalNAc and other glycans by isotopic tagging using C1GalT1 and UDP-Gal(^13^C_6_), and map Tn-glycosylation sites using OpeRATOR and LC-MS/MS.

**Figure 2.**
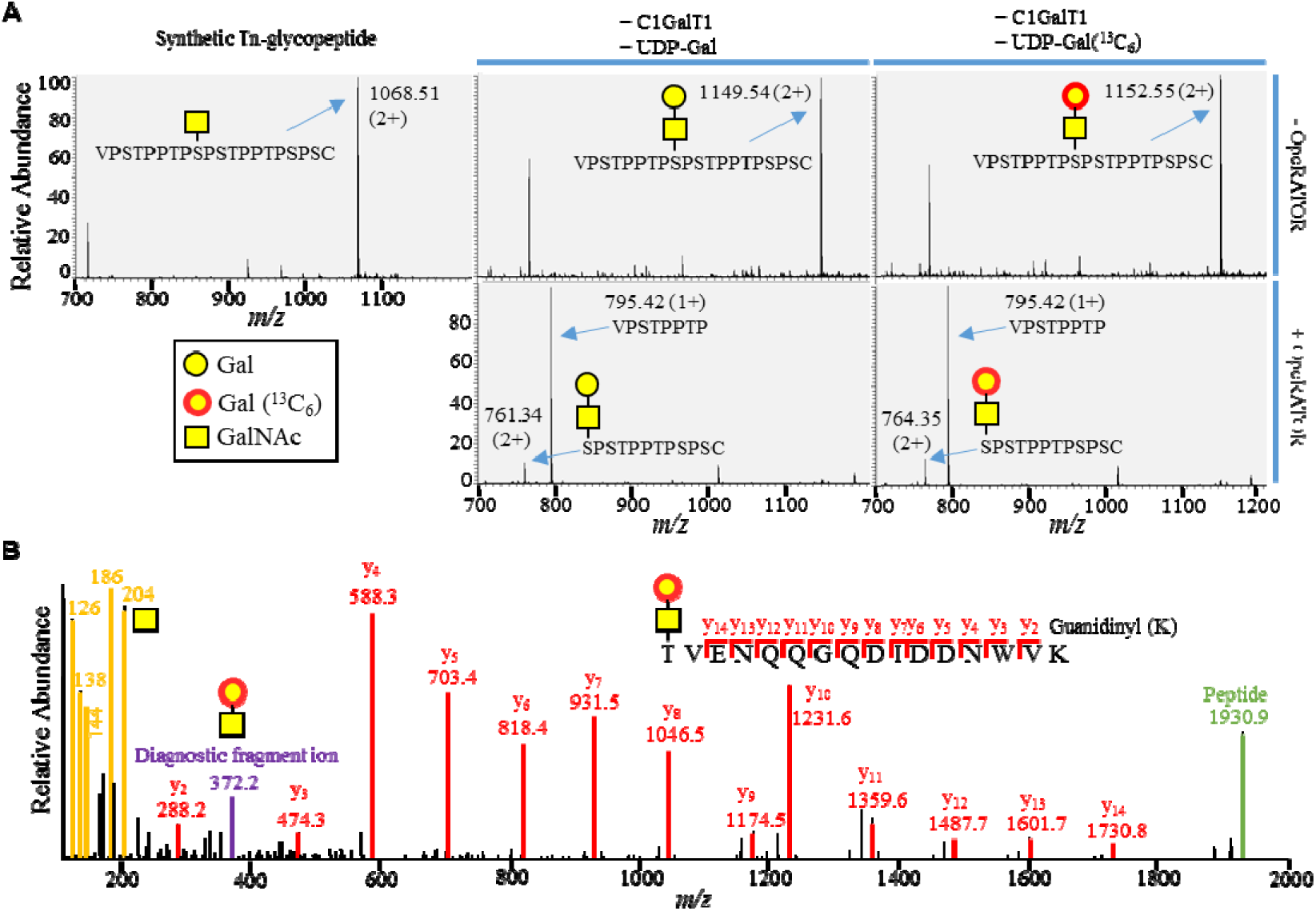
Mapping Tn-glycosylation sites by integrating Tn-engineering and OpeRATOR digestion. A. OpeRATOR digestion of Gal- and Gal(^13^C_6_)-Tn-glycopeptide after Tn was tagged using C1GalT1 with UDP-Gal or UDP-Gal(^13^C_6_). Top left panel: the synthetic Tn-glycopeptide before treatments. Top middle panel: conversion of Tn to Gal-Tn using C1GalT1 and UDP-Gal. Bottom middle panel: OpeRATOR digestion of the Gal-Tn-glycopeptide generated in the top middle panel produced site-containing glycopeptide S(Gal-Tn)PSTPPTPSPSC-NH2 and peptide VPSTPPTP. Top right panel: conversion of Tn to Gal(^13^C_6_)-Tn using C1GalT1 and UDP-Gal(^13^C_6_). Bottom right panel: OpeRATOR digestion of the Gal(^13^C_6_)-Tn-glycopeptide engineered in the top right panel yielded site-containing glycopeptide S(Gal(^13^C_6_)-Tn)PSTPPTPSPSC-NH2 and peptide VPSTPPTP. B. HCD-MS2 spectrum of site-containing Gal(^13^C_6_)-Tn-glycopeptide identified in Jurkat cells. A diagnostic oxonium ion at 372 *m/z* corresponding to fragmentation ion of Gal(^13^C_6_)-Tn was colored in purple.

### Mapping site-specific Tn-glycoproteome in Jurkat cells

Jurkat cells were analyzed to evaluate the performance of EXoO-Tn. With 1% FDR, 3,172 peptide-spectrum match (PSM) were assigned to 1,078 unique site-containing Gal(^13^C_6_)-Tn-glycopeptides that contained 1,011 unique peptide sequences (Fig. 3 and Supplementary Table 1). From the peptide sequence, we mapped 947 Gal(^13^C_6_)-Tn-glycosylation sites from 480 glycoproteins (Fig. 3 and Supplementary Table 1). The diagnostic oxonium ion at 372 *m/z* was detected in 96.4% of the assigned MS/MS spectra with an overall intensity being ten-fold lower than that at 204 *m/z* (Fig. 4A and Supplementary Table 1). The detection of oxonium ion at 372 *m/z* in the assigned MS2 spectra supported the presence of Gal(^13^C_6_)-Tn in the identified glycopeptides (Supplementary Table 1). It was observed that, among the assigned PSMs, approximately 89.2% glycopeptides were modified by a single Gal(^13^C_6_)-Tn composition while approximately 9.5 and 1.3% PSMs were modified by two or three Gal(^13^C_6_)-Tn compositions, respectively (Supplementary Table 1).

**Figure 3.**
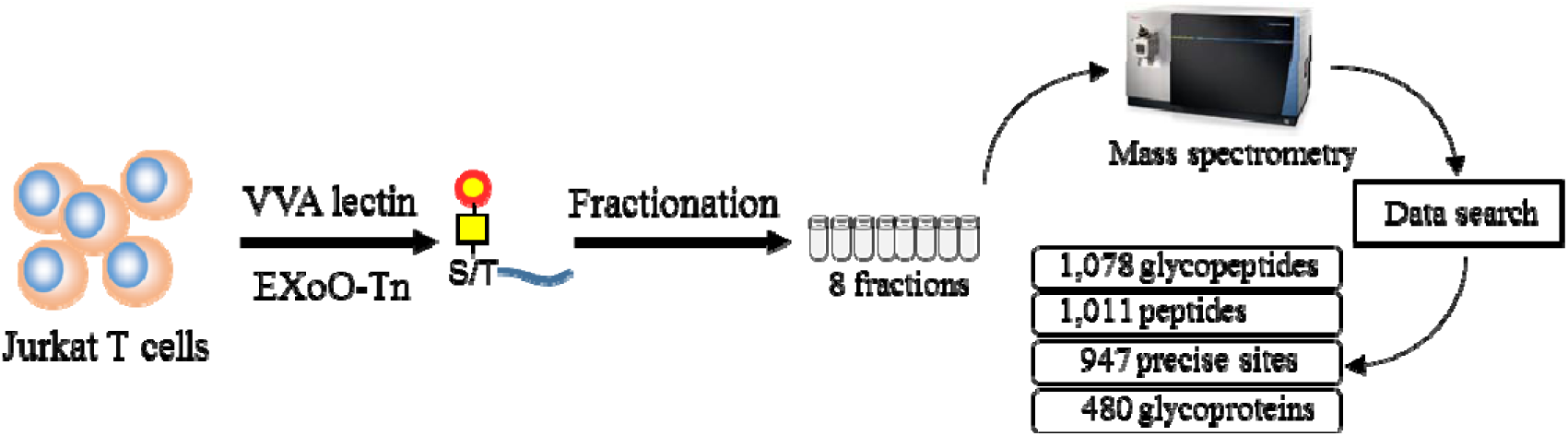
A Schematic workflow for identification of site-specific Tn-glycoproteome in Jurkat cells.

**Figure 4.**
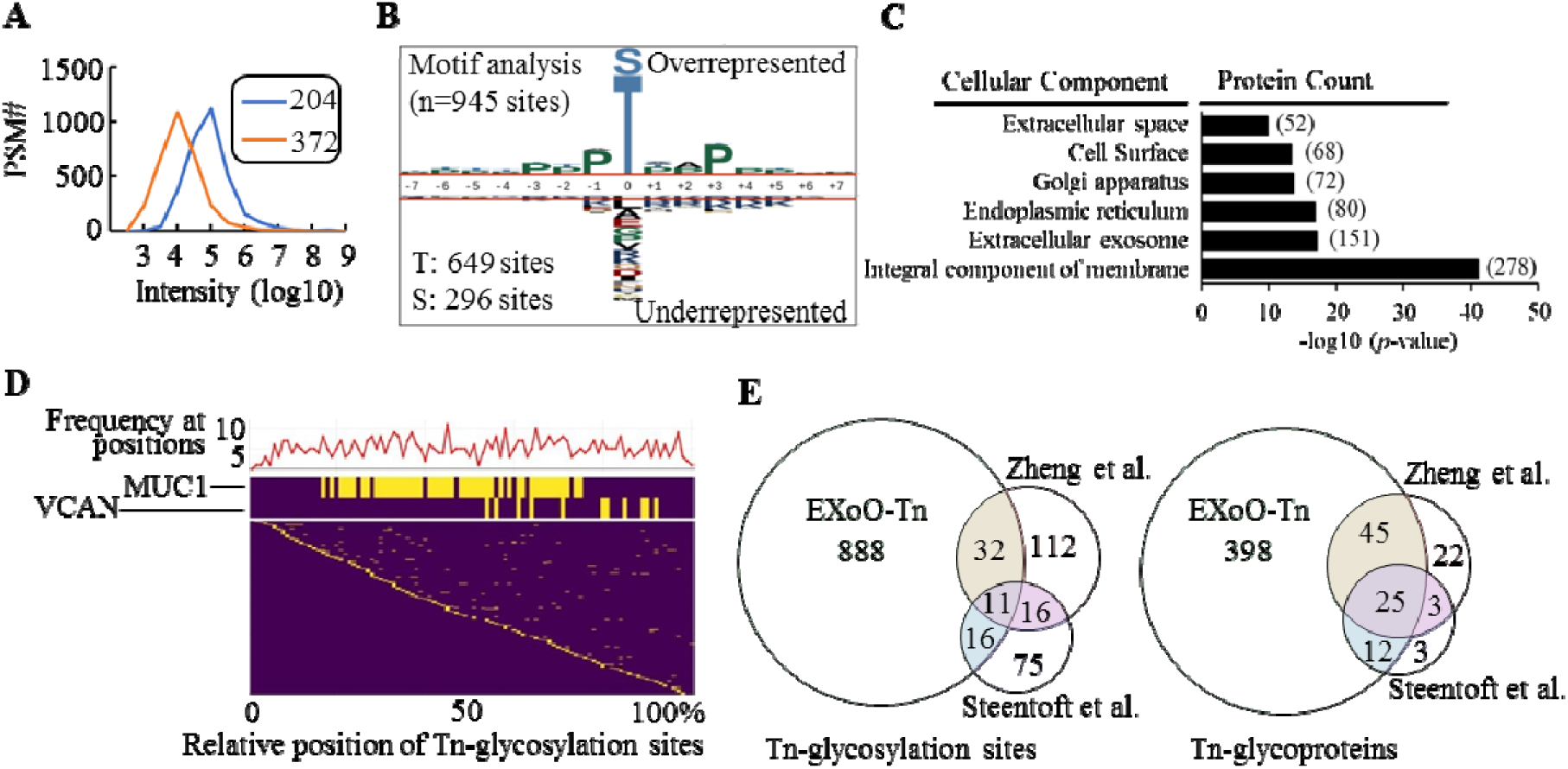
Characteristics of site-specific Tn-glycoproteome in Jurkat cells. A. The overall intensity of oxonium ions at 204 and 372 *m/z* in the assigned PSMs. The overall intensity of oxonium ion at 372 *m/z* was 10-fold less than that of 204 *m/z*. B. Motif analysis revealed the conserved motif of Tn-glycosylation sites. C. GO analysis revealed cellular components for Tn-glycoproteome. D. Analysis of the relative position of Tn-glycosylation sites in protein sequences revealed that the frequency of Tn-glycosylation distributed evenly across protein sequences with lower frequency at protein termini. E. Comparison of O-linked glycosylation sites and glycoproteins identified in this and other studies ^19, 20^.

### Characterization of the site-specific Tn-glycoproteome in Jurkat cells

Analysis of the glycosylation sites showed that Thr and Ser accounted for approximately 68.7% and 31.3%, respectively. Motif analysis of ±7 amino acids surrounding 946 glycosylation sites found an overrepresentation of Pro residues at the +3 and −1 position (Fig. 4B). Two glycosylation sites residing close to the protein N-termini were not used in the motif analysis. Gene Ontology (GO) analysis of the identified glycoproteins found that integral component of membrane, extracellular exosome, endoplasmic reticulum (ER), Golgi apparatus, cell surface, and extracellular space were enriched for cellular component suggesting the presence of the identified glycoproteins in the secretory pathway and on the cell surface (Fig. 4C). Next, the relative position of the glycosylation sites in protein sequence was plotted and showed that proteins MUC1 and versican core protein (VCAN) had the highest number of glycosylation sites reaching 48 and 11, respectively (Fig. 4D middle panel). Besides, it was observed that the frequency of the glycosylation site was relatively even across protein sequences with lower frequency at protein termini (Fig. 4D top and bottom panels). Comparison of site-specific Tn-glycoproteome identified by EXoO-Tn to two other methods ^19, 20^ (Supplementary Table 2 and 3) revealed that 888 Tn-glycosylation sites from 398 glycoproteins were exclusively identified using EXoO-Tn (Fig. 4E). Analysis of Jurkat cells established the effectiveness of EXoO-Tn to map the site-specific Tn-glycoproteome in the complex sample.

## Discussion

A new technology EXoO-Tn has been developed for large-scale mapping Tn-glycosylation sites in the complex sample. EXoO-Tn has several advantages including (i) large-scale mapping of Tn-glycosylation sites in the complex sample; (ii) a tagging strategy for distinguishing engineered Tn from endogenous Gal-GalNAc and other glycans; (iii) concurrent tagging of Tn and release of site-containing Tn-glycopeptides from solid-phase in a one-pot fashion; (iv) applicable to analyze mucin-type O-linked glycoproteins; (v) no need of ETD for site localization.

C1GalT1 is a natural enzyme with specificity for extending O-GalNAc to core 1 Gal-GalNAc structure. OpeRATOR enzyme is utilized by bacteria to digest mucin glycoproteins in the gut with a specificity at N-termini of Gal-GalNAc occupied Ser/Thr residues. The two enzymes work synergistically to render EXoO-Tn the specificity for mapping Tn-glycosylation sites. It is meritorious that Tn is tagged to have a unique mass and generate a diagnostic oxonium ion in the MS2 spectrum. The unique mass tag and diagnostic oxonium ion are useful to improve the confidence of identification. The use of solid-phase allows extensive washes that are essential to remove other peptides and contaminants while enables further enrichment of site-containing glycopeptides for LC-MS/MS analysis.

We mapped 947 Tn-glycosylation sites from almost 500 glycoproteins, a substantially large number of site-specific Tn-glycoproteome, which demonstrated the effectiveness of EXoO-Tn and supported that a large number of O-linked glycosylation sites could be mapped in cells. Some site-containing Tn-glycopeptides may be too long or too short to be detected using EXoO-Tn with trypsin digestion. Digestion of proteins using proteases with different specificities may further increase the identification number of glycosylation sites in EXoO-Tn methodology. Also, the identification of glycopeptides with two or three Gal(^13^C_6_)-Tn compositions suggests many more glycosylation sites in the peptide sequences supporting an even larger number of Tn-glycosylation sites in Jurkat cells. Characterization of glycosylation sites and glycoproteins identified in Jurkat cells revealed conserved features of protein O-linked glycosylation, including consensus motif, cellular localization, and distribution of the relative position of glycosylation sites across the protein sequences, a reminiscence of that seen in human kidney, serum, and T cells in the previous study ^21^. Given that Tn is prevalent in cancers and other diseases, EXoO-Tn is anticipated to have broad translational and clinical utilities.

## Material and Methods

### Tagging of Tn and mapping its glycosylation site using synthetic Tn-glycopeptide

Synthetic Tn-glycopeptide VPSTPPTPS(α-GalNAc)PSTPPTPSPSC-NH2 IgA1 hinge peptide was purchased from Susses Research. In the workflow with sequential enzymatic treatments, five µg of glycopeptide in 50 mM Tris-HCl pH 7.4 was mixed with one µg recombinant human C1GalT1/C1GalT1C1 protein (R&D Systems, NM) in the presence of either 0.5 mM UDP-Gal (Sigma-Aldrich) or 0.5 mM UDP-Gal^13^C_6_ (Omicron Biochemicals, lnc., IN) at 37°C for 16 hours. After incubation, half of each sample was subjected to digestion using five units of OpeRATOR (Genovis Inc, Cambridge, MA) at 37°C for 16 hours. The glycopeptides were desalted using C18 ZipTip (Millipore Sigma), dried using speed-vac, and resuspended in 0.1% TFA. In the concurrent one-pot enzymatic treatment that was used in all experiments described below, enzymes including C1GalT1/C1GalT1C1, OpeRATOR, and substrate i.e. UDP-Gal or UDP-Gal^13^C_6_ were added at the same time using the amount as described in the above sequential enzymatic workflow and incubated at 37°C for 16 hours before C18 desalting and LC-MS/MS analysis.

### Extraction of site-containing Tn-glycopeptides from Jurkat cells

Jurkat Clone E6-1 (NIH AIDS Reagent Program) were cultured and expanded in RPMI 1640 supplemented with 10% fetal bovine serum (FBS), 100 units of penicillin, and 100 µg of streptomycin. The cells were collected, washed three times in the ice-cold PBS and lysed in 8 M urea/500 mM ammonia bicarbonate. The cell lyse was sonicated and centrifuged at 16,000 g to remove particles. Protein concentration was determined using a protein BCA assay. Twenty milligrams of proteins were reduced in 5 mM DTT at 37°C for 1 hour and alkylated in 10 mM iodoacetamide at room temperature (RT) for 40 min in the dark. The samples were then diluted five-fold using 100 mM ammonia bicarbonate buffer. Trypsin was added to the samples with an enzyme/protein ratio of 1/40 w/w. After incubation at 37°C for 16 hours, lysine residues were guanidination-modified, and peptides were desalted using C18 cartridges (Waters, Milford, MA), as described in the previous study ^21^. The peptides were dried using speed-vac, resuspended in PBS with α2-3,6,8 neuraminidase (New England Biolabs, Ipswich, MA), and incubated at 37°C for 16 hours. Four-hundred microliters agarose bound Vicia Villosa Lectin (VVA) (50% slurry, Vector Laboratories, Burlingame, CA) were washed twice using water, added to peptides and incubated at RT for 16 hours with rotation. The VVA agarose was gently washed with 1X PBS for three times. Bound glycopeptides were eluted using 4 M urea/100 mM Tris-HCl pH 7.4/400mM GalNAc (Sigma-Aldrich) at RT for 30 min with shaking. The eluted glycopeptides were desalted using C18 cartridge and conjugated to AminoLink resin (Pierce, Rockford, IL) as described previously ^21^. Briefly, the pH of C18 elute containing glycopeptides was neutralized to approximately pH 7 using two volume of 10X PBS. The solution was mixed with resin (100 µg peptide/100 µl resin, 50% slurry) and 50 mM sodium cyanoborohydride (NaCNBH_3_) at RT for a minimal of 4 hours or overnight with rotation. Unreacted groups on resin were blocked using 1M Tris-HCl buffer (pH7.4) with 50 mM NaCNBH_3_ at RT for 30 min with rotation. The resin was sequentially washed using 50% ACN, 1.5 M NaCl, and 50 mM Tris-HCl buffer (pH 7.4). To tag and release Tn-glycopeptides, a solution (50 µl) containing 10 µg of C1GalT1/C1GalT1C1, 0.5 mM UDP-Gal^13^C_6_, and 2000 units of OpeRATOR was added to the resin and incubated at 37°C for 16 hours. The released glycopeptides in the solution were collected twice using 400 µl of 50 mM Tris-HCl buffer (pH 7.4). Glycopeptides in the collected solution were combined, desalted using C18 cartridge, dried using speed-vac, and resuspended in 0.1% TFA. The peptides were fractionated using HPLC and concatenated to eight fractions before LC-MS/MS analysis.

### LC-MS/MS analysis

One microgram of glycopeptides was analyzed on a Fusion Lumos mass spectrometer with an EASY-nLC 1200 system or an LTQ Orbitrap Velos mass spectrometer (Thermo Fisher Scientific, Bremen, Germany). The mobile phase flow rate was 0.2 μl/min with 0.1% FA/3% acetonitrile in water (A) and 0.1% FA/90% acetonitrile (B). The gradient profile was set as follows: 6% B for 1 min, 6-30% B for 84 min, 30-60% B for 9 min, 60-90% B for 1 min, 90% B for 5 min and equilibrated in 50% B, flow rate was 0.5 μL/min for 10 min. MS analysis was performed using a spray voltage of 1.8 kV. Spectra (AGC target 4 × 10^5^ and maximum injection time 50 ms) were collected from 350 to 1800 m/z at a resolution of 60 K followed by data-dependent HCD MS/MS (at a resolution of 50 K, collision energy 36, AGC target of 2 × 10^5^ and maximum IT 250 ms) of the 15 most abundant ions using an isolation window of 0.7 m/z. Include charge state was 2-6. The fixed first mass was 110 m/z. Dynamic exclusion duration was 45 s.

### Database search of site-containing Tn-glycopeptides

A UniProt human protein database (71,326 entries, downloaded October 19, 2017) was used to generate a peptide database with 26,067,074 non-redundant peptide entries using the method as described in the previous study ^21^. Briefly, a randomized decoy database using The Trans-Proteomic Pipeline (TPP) ^22^ was generated and concatenated with the target database. The concatenated database was digested with trypsin and then OpeRATOR in silico. Peptides with Ser or Thr residues and lengths from 6 to 46 amino acids were used. SEQUEST in Proteome Discoverer 2.2 (Thermo Fisher Scientific) was used to search with variable modification: oxidation (M), Gal^13^C_6_(1)HexNAc(1) (S/T), Hex(1)HexNAc(1) (S/T) and HexNAc (S/T) and static modification: carbamidomethylation (C) and guanidination (K). FDR was set at 1% using Percolator. Only MS/MS scans with oxonium ion at 204, and two of the other oxonium ions were kept. Assignments with XCorr score below one were removed. MS/MS spectra were manually studied and inspected using spectral viewer in Proteome Discoverer to identify the spectral feature and ensure the confidence of identification.

### Bioinformatics

Software pLogo was used to reveal motif for Tn-glycosylation sites ^23^ surrounding by 15 amino acids in length with the central amino acids being the sites. The Database for Annotation, Visualization and Integrated Discovery (DAVID) and UniProt (http://www.uniprot.org) were used for Gene Ontology (GO) analysis ^24^. Python (version 2.7) is used to analyze the data and generate the figures, including the relative position of Tn-glycosylation sites in protein sequence, radar charts, unsupervised hierarchical clustering, and box plot.

## Data Availability

The LC-MS/MS data have been deposited to the PRIDE partner repository ^25^ with the dataset identifier: project accession: PXD014390

Reviewer account details:

Username

Password: tZVBNHhu

## Acknowledgment

We acknowledge Dr. Kyung-Cho Cho for maintenance of Mass Spectrometer. This work was supported by the National Cancer Institute, the Early Detection Research Network (EDRN, U01CA152813), the Clinical Proteomic Tumor Analysis Consortium (CPTAC, U24CA210985), the National Institute of Allergy and Infectious Diseases (R21AI122382), and by amfAR, The Foundation for AIDS Research on Bringing Bioengineers to Cure HIV (Grant amfAR 109551-61-RGRL). The following reagent was obtained through the NIH AIDS Reagent Program, Division of AIDS, NIAID, NIH: Jurkat Clone E6-1 from Dr. Arthur Weiss (cat# 177) ^26^.

## Contributions

W.Y. and H.Z. conceived the concept and wrote the manuscript; W.Y., A.S., and Y.X. conducted experiments and data analysis; M.A performed programming, advanced data analysis, and bioinformatics.

## References

1. Julien, S., Videira, P.A. & Delannoy, P. Sialyl-tn in cancer: (how) did we miss the target? Biomolecules 2, 435–466 (2012).

2. Munkley, J. The Role of Sialyl-Tn in Cancer. International journal of molecular sciences 17, 275 (2016).

3. Ju, T. et al. Tn and sialyl-Tn antigens, aberrant O-glycomics as human disease markers. Proteomics. Clinical applications 7, 618–631 (2013).

4. Kudelka, M.R., Ju, T., Heimburg-Molinaro, J. & Cummings, R.D. Simple sugars to complex disease--mucin-type O-glycans in cancer. Advances in cancer research 126, 53–135 (2015).

5. Slovin, S.F. et al. Fully synthetic carbohydrate-based vaccines in biochemically relapsed prostate cancer: clinical trial results with alpha-N-acetylgalactosamine-O-serine/threonine conjugate vaccine. Journal of clinical oncology: official journal of the American Society of Clinical Oncology 21, 4292–4298 (2003).

6. Itzkowitz, S.H., Bloom, E.J., Lau, T.S. & Kim, Y.S. Mucin associated Tn and sialosyl-Tn antigen expression in colorectal polyps. Gut 33, 518–523 (1992).

7. Inoue, M., Ton, S.M., Ogawa, H. & Tanizawa, O. Expression of Tn and sialyl-Tn antigens in tumor tissues of the ovary. American journal of clinical pathology 96, 711–716 (1991).

8. Wei, H. et al. Glycoprotein screening in colorectal cancer based on differentially expressed Tn antigen. Oncology reports 36, 1313–1324 (2016).

9. Nakagoe, T. et al. Prognostic value of circulating sialyl Tn antigen in colorectal cancer patients. Anticancer research 20, 3863–3869 (2000).

10. Tsuchiya, A. et al. Prognostic Relevance of Tn Expression in Breast Cancer. Breast cancer 6, 175–180 (1999).

11. Ohno, S. et al. Expression of Tn and sialyl-Tn antigens in endometrial cancer: its relationship with tumor-produced cyclooxygenase-2, tumor–infiltrated lymphocytes and patient prognosis. Anticancer research 26, 4047–4053 (2006).

12. Posey, A.D., Jr. et al Engineered CAR T Cells Targeting the Cancer-Associated Tn-Glycoform of the Membrane Mucin MUC1 Control Adenocarcinoma. Immunity 44, 1444–1454 (2016).

13. Wilkie, S. et al. Retargeting of human T cells to tumor-associated MUC1: the evolution of a chimeric antigen receptor. Journal of immunology 180, 4901–4909 (2008).

14. Maher, J. et al. Targeting of Tumor-Associated Glycoforms of MUC1 with CAR T Cells. Immunity 45, 945–946 (2016).

15. Ju, T. et al. Human tumor antigens Tn and sialyl Tn arise from mutations in Cosmc. Cancer research 68, 1636–1646 (2008).

16. Hofmann, B.T. et al. COSMC knockdown mediated aberrant O-glycosylation promotes oncogenic properties in pancreatic cancer. Molecular cancer 14, 109 (2015).

17. Moran, S. & Cattran, D.C. IgA nephropathy: un update. Minerva medica (2019).

18. Berger, J. & Hinglais, N. [Intercapillary deposits of IgA-IgG]. Journal d’urologie et de nephrologie 74, 694–695 (1968).

19. Steentoft, C. et al. Mining the O-glycoproteome using zinc-finger nuclease-glycoengineered SimpleCell lines. Nature methods 8, 977–982 (2011).

20. Zheng, J., Xiao, H. & Wu, R. Specific Identification of Glycoproteins Bearing the Tn Antigen in Human Cells. Angewandte Chemie 56, 7107–7111 (2017).

21. Yang, W., Ao, M., Hu, Y., Li, Q.K. & Zhang, H. Mapping the O-glycoproteome using site-specific extraction of O-linked glycopeptides (EXoO). Mol Syst Biol 14, e8486 (2018).

22. Deutsch, E.W. et al. Trans-Proteomic Pipeline, a standardized data processing pipeline for large-scale reproducible proteomics informatics. Proteomics. Clinical applications 9, 745–754 (2015).

23. O’Shea, J.P. et al. pLogo: a probabilistic approach to visualizing sequence motifs. Nature methods 10, 1211–1212 (2013).

24. Huang da, W., Sherman, B.T. & Lempicki, R.A. Systematic and integrative analysis of large gene lists using DAVID bioinformatics resources. Nat Protoc 4, 44–57 (2009).

25. Vizcaino, J.A. et al. 2016 update of the PRIDE database and its related tools. Nucleic acids research 44, D447–456 (2016).

26. Weiss, A., Wiskocil, R.L. & Stobo, J.D. The role of T3 surface molecules in the activation of human T cells: a two-stimulus requirement for IL 2 production reflects events occurring at a pre-translational level. Journal of immunology 133, 123–128 (1984).

